# The most common epilepsy-causing mutation in EEF1A2 (E122K) perturbs the translation of specific transcripts but not the rate of global protein synthesis

**DOI:** 10.64898/2026.07.08.737232

**Authors:** Cavan Bennett Ness, Manuela Rizzi, Heather Love, Nika Balkic, Grant Marshall, Alex von Kriegsheim, Emily Osterweil, Catherine M. Abbott

**Affiliations:** Institute of Genetics and Cancer, University of Edinburgh, Western General Hospital, Edinburgh EH4 2XU, UK; Simons Initiative for the Developing Brain, University of Edinburgh, Edinburgh, UK; Centre for Discovery Brain Sciences, University of Edinburgh, Edinburgh, UK

## Abstract

Heterozygous *de novo* missense mutations in the *EEF1A2* gene encoding translation elongation factor eEF1A2 result in neurodevelopmental disorders, typically characterised by early onset epilepsy and intellectual disability (ID). The E122K mutation is the most commonly reported missense mutation and is amongst the more severe in terms of epilepsy and ID. Here we made use of a recently developed mouse model which recapitulates the E122K mutation to examine how mutations in *EEF1A2* might disrupt neuronal gene expression. Primary neurons from mutant mice and transfected HEK293T cells were used to examine effects on global protein synthesis. In contrast to previous reports, we were unable to detect a change in global protein synthesis using either of two different assay systems. TRAP-seq and mass spectrometry were then employed to study the effects of the mutation on the translatome and proteome respectively. These analyses revealed perturbation of expression of a subset of genes, with a slight skew towards downregulation, particularly for longer transcripts. Further analysis indicated a down regulation of proteins involved in synaptic function in both the translatomic and proteomic datasets.

## Introduction

Heterozygous *de novo* mutations in the *EEF1A2* gene result in *EEF1A2*-Related Neurodevelopmental Disorder, which is characterised by early onset epilepsy, autism, and intellectual disability (ID), sometimes with a degenerative course [1]. More than 50 independent missense mutations have been identified in the gene [2], with an incidence rate of 2.92 per 100,000 births [3]. Around half of these missense mutations have only been reported in a single individual, but for the more common mutations, like G70S, E122K, E124K and R266W, genotype-phenotype correlations are emerging, with some mutations consistently being associated with a more severe phenotype than others. The E122K mutation is the most commonly reported of all missense mutations and is generally associated with intractable seizures and moderate to severe ID [1, 4, 5].

Although there has been some debate about whether these mutations represent a gain or loss of function [1], the mutational profile of almost exclusively missense mutations is suggestive of a gain of function or dominant negative mechanism since haploinsufficiency for *EEF1A2* is compatible with life, and generally associated with milder phenotypes [6, 7]. Direct evidence for the gain of function argument comes from mouse models of the E122K and D252H mutations where in each case the phenotype is more severe in animals heterozygous or homozygous for missense mutations than that seen in the equivalent nulls [8, 9].

The *EEF1A2* gene encodes a tissue-specific form of eukaryotic elongation factor 1A (eEF1A), the central functional component of the multi-subunit eEF1 complex. eEF1A is responsible for the delivery of aminoacyl tRNAs to the A-site of the ribosome to extend the nascent polypeptide chain during protein synthesis. Vertebrates have two independently encoded eEF1A isoforms, eEF1A1 and eEF1A2, which are 92% identical at the amino acid level but expressed in a reciprocal, tissue-specific manner [10–12]. Throughout embryogenesis and early development, eEF1A1 is expressed ubiquitously, whereas eEF1A2 is expressed only in brain, cardiac and skeletal muscle, replacing eEF1A1 in muscle once development is complete. Early reports suggested that as eEF1A2 expression increases, eEF1A1 expression becomes downregulated to the point of being undetectable in brain [13]. However, whilst neurons within the mature central nervous system express eEF1A2, glia exclusively express eEF1A1 [14] [15], complicating analysis in this tissue. More recent work using a mouse line in which a V5 epitope tag was expressed from the endogenous eEF1A2 locus has shown that eEF1A2 expression can be detected as early as E11.5 and that, whilst eEF1A2 is indeed exclusively expressed in the neuronal soma, eEF1A1 expression is in fact retained in axons. There is thus not only a developmental switch from eEF1A1 to eEF1A2, but also a postnatal relocation of eEF1A1 expression within neurons resulting in complete compartmentalisation of the two proteins [16].

This entirely mutually exclusive expression pattern for two near-identical proteins of course raises the issue of functional equivalence of eEF1A1 and eEF1A2, and whether any differences relate to the canonical function of the proteins in translation, or to non-canonical functions. Many of these non-canonical functions have been attributed in the literature simply to eEF1A (without distinguishing between variants) ever since it was first noted that eEF1A is in excess over other translation elongation factors [17]. The most notable of these functions is cytoskeletal organisation [18], and a role for eEF1A in the transport of mRNAs to neuronal synapses in RNA transport granules has been reported [19].

Some isoform-specific functions have also been identified. eEF1A1 has been shown to promote the expression of HSP70 through interaction with heat shock factor 1 (HSF1), whilst eEF1A2 was not sufficient to induce a response [20]. Phosphorylation of eEF1A2 on ser205 and ser358 (residues which are both alanine in eEF1A1) by c-Jun N-terminal kinase (JNK) was found to promote proteasomal degradation in response to cellular stress [21]. Similarly, although both isoforms have been found to bind the actin cytoskeleton, *in vitro* studies have shown the isoforms have different actin bundling properties [22–25] and Mendoza et al further showed that phosphorylation of four eEF1A2-specific serine residues (including ser358) coordinates the activity of eEF1A2 in protein synthesis and actin remodelling [23]. [26–28]

A central question is how pathogenic missense mutations in eEF1A2 give rise to neurodevelopmental disorders, and whether they impact on canonical or non-canonical functions, or both. There is strong evidence for dysregulation of translation as an underlying mechanism in neurodevelopmental disorders like tuberous sclerosis and Fragile X syndrome, so it is reasonable to assume a similar perturbation exists in *EEF1A2*-Related Neurodevelopmental Disorder. In support of this, Mohamed and Klann [29] recently demonstrated that three disease-causing eEF1A2 mutations, G70S, E122K, and D252H, decrease *de novo* protein synthesis in HEK293T cells and mouse cortical neurons and alter neuronal morphology, suggesting that *EEF1A2-*related disorders may result from a loss of protein synthesis. The mutations also showed increased tRNA binding and decreased actin binding, with implications for mRNA transport, local translation, and axonal growth.

In this study, we made use of a recently developed mouse model which recapitulates the E122K mutation, the most common cause of *EEF1A2*-Related Neurodevelopmental Disorder, [30] to re-examine the question of how mutations disrupt neuronal gene expression. Mice heterozygous for the mutation develop lifelong body mass deficits, transient early motor delays, and electrographic seizures. Consistent with results from mice with other biallelic *Eef1a2* mutations [8, 31], phenotypic measurements show an earlier onset of neurological dysfunction in mice homozygous for the E122K mutation in comparison with homozygous null mice, consistent with a toxic gain of function. We used both primary neurons from the mice and transfected cells to examine effects on global protein synthesis. In contrast to previous reports, we were unable to detect a change in total protein synthesis using either of two methods. As we also found no evidence for proteotoxic/ribotoxic stress, we went on to use TRAP-seq and mass spectrometry to study the effects of the mutation on the translatome and proteome respectively. We observed the perturbation of expression of a subset of genes, with a slight skew towards downregulation. Overall, translatome and proteome profiles reveal a poor overlap between mRNA and protein expression, though GSEA revealed the convergent downregulation of genes relating to synaptic function.

## Results

### Effects of missense mutations on protein synthesis

We initially carried out protein synthesis assays in primary neurons derived from a mouse line carrying the E122K mutation in the endogenous *Eef1a2* gene, the most common pathogenic variant seen in humans (referred to here as E122K mice). We wanted to ask if the reduction in protein synthesis previously reported in HEK cells and primary neurons transfected with an E122K expression construct [29] was also observed in the absence of transfection. We used the SUnSET assay, in which puromycin is incorporated into newly synthesised proteins then detected using an anti-puromycin antibody.[32] We found no evidence of a decrease in protein synthesis in DIV14 neurons from E122K mice, even when homozygous mutants were compared to those from wild-type (WT) littermates (Figure 1A). Although some variability was observed between samples, puromycin incorporation was not significantly altered by genotype. To confirm this result with an independent approach, we then measured protein synthesis in primary neurons from the same mouse line using an assay based on incorporation of l-azidohomoalanine (AHA), which is subsequently fluorescently labelled with click chemistry. Consistent with the SUnSET assay, this also revealed no significant difference between genotypes (Figure 1C), with a slight but non-significant increase in the mutants. AHA incorporation decreased significantly however in the presence of cycloheximide, an inhibitor of protein synthesis.

**Figure 1:**
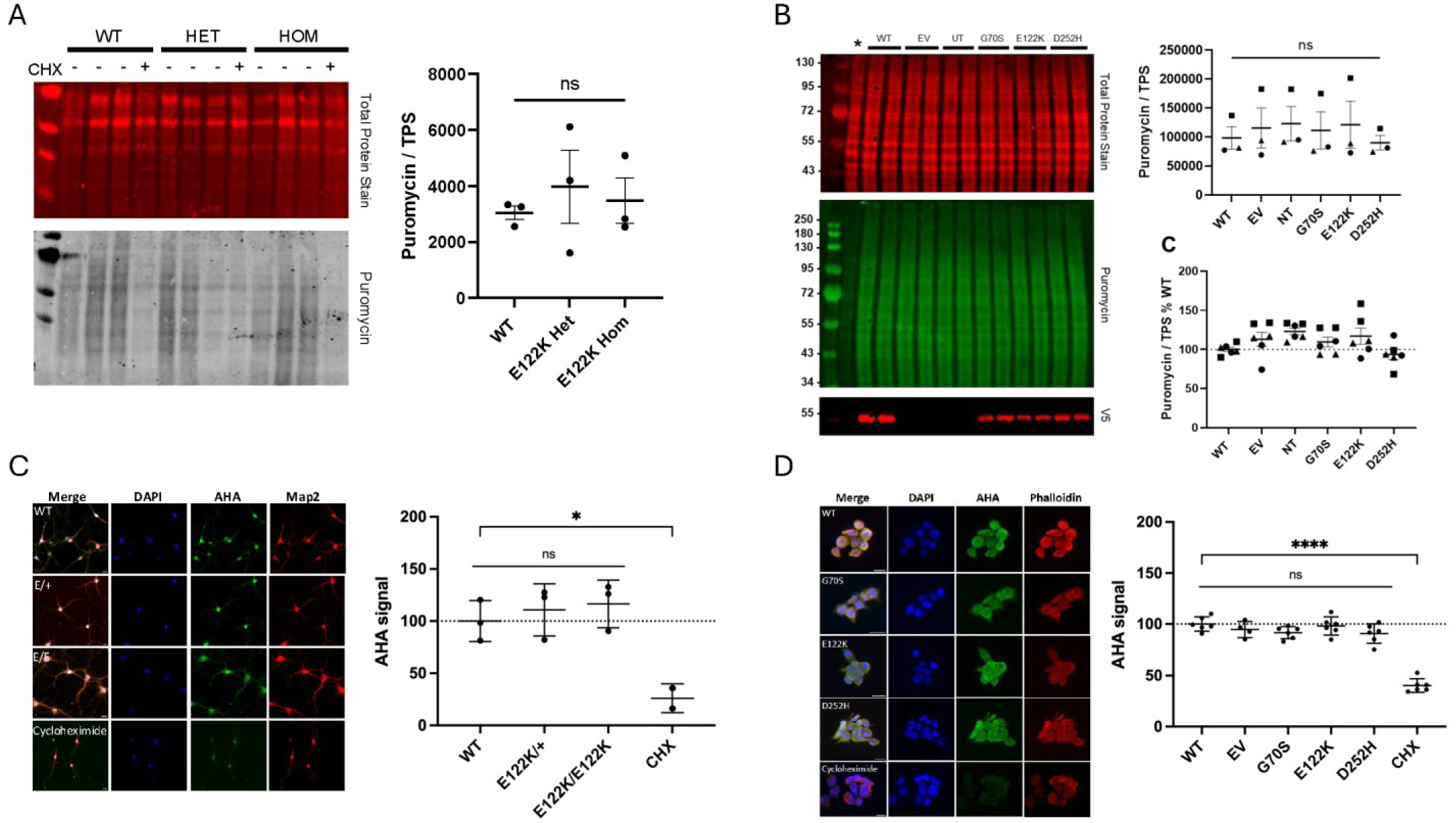
Protein synthesis is not significantly altered by pathogenic eEF1A2 variants in HEK293T cells or mouse primary neurons. A) *SUnSET assay in DIV14 hippocampal cortical neuronal co-cultures from eEF1A2 WT, heterozygous, and homozygous E122K mice. Neurons were exposed to 1 µg/ml puromycin for 30 minutes, then analysed by western blotting to measure incorporation. CHX + lanes show control cells treated with 50 µg/ml cycloheximide for 30 minutes in addition to puromycin. Blots were probed for puromycin and quantified in Image Studio Lite, with puromycin incorporation displayed relative to total protein. A one-way ANOVA showed no significant difference in puromycin incorporation between the three genotypes (F (2,6) = 0.266, p = 0.775)*. B) *SUnSET assay in HEK293T cells transiently transfected with V5-tagged empty vector (EV), WT, G70S, E122K, or D252H EEF1A2 cDNA within a pcDNA3.1 vector, or left untransfected (UT). Cells were treated with puromycin and cycloheximide as described above, where * indicates puromycin + cycloheximide control. Representative blots shown were probed with anti-V5 (red) and anti-puromycin (green) antibodies. Quantification (right) shows puromycin incorporation normalised to total protein (top), with each point representing an average of two lanes from a replicate, and each replicate indicated by a different point shape. A repeated measures one-way ANOVA found no significant difference in incorporated puromycin signal between any condition after transfection of variant eEF1A2 (F (2, 10) = 1.578, P = 0.252). The lower panel shows quantified eEF1A2 signal for every individual lane, normalised to the total protein stain and then calculated as a proportion of the control condition for visualisation*. C) *Click-iT® AHA assay in mouse primary neuronal cultures. Neurons were cultured as in A., treated with 50 µM Click-iT® L-azidohomoalanine (AHA) reagent for 30 minutes, then processed for fluorescent imaging. Images were acquired using an EVOS fluorescence microscope and analysed in QuPath. Representative images of neurons showing AHA incorporation in neurons alongside a neuronal marker MAP2. Scale bar: 20 µm. Quantification was performed in QuPath and shows mean AHA signal per cell body normalised to WT. A one-way ANOVA showed no significant difference in normalised AHA levels between the three genotypes (F (2,6) = 0.411, p = 0.68))*. D) *Click-iT® AHA assay performed as described above in C., carried out in HEK293T cells transfected with WT or mutant eEF1A2 constructs as in B. Representative epifluorescence images of cells treated with AHA reagent. Scale bar: 20 µm. Quantification shows mean AHA signal per cell expressed relative to WT. A one-way ANOVA revealed no significant difference between conditions (F(3, 20) = 2.039, p = 0.137)*.

In view of this surprising discrepancy with the literature, we then repeated the same assays in transfected HEK cells, which do not normally express endogenous eEF1A2 but rather rely on eEF1A1. Importantly, we also included untransfected cells as a control, as the results obtained by Mohamed and Klann [29], where protein synthesis was impaired in cells transfected with mutant forms of eEF1A2 compared with those transfected with WT eEF1A2, could be interpreted in two ways. Firstly, there could be a dominant negative effect of mutant eEF1A2 on the function of eEF1A1, or alternatively, transfected eEF1A2 could enhance protein synthesis in the cells, but to a lesser extent for mutant than WT eEF1A2; the addition of untransfected cells helps to distinguish between these possibilities. As with the primary neurons, we saw no difference in *de novo* protein synthesis in HEK cells transfected with any of three pathogenic eEF1A2 variants (G70S, E122K and D252H), using either puromycin or AHA incorporation assays (Figure 1B, 1D). We also saw no difference between cells transfected with WT eEF1A2 or empty vector, suggesting that the addition of eEF1A2 has no detectable effect on the activity of endogenous eEF1A1. Overall, we were unable to observe any measurable effect of mutant eEF1A2 on protein synthesis in cells expressing eEF1A1, regardless of origin.

### The E122K mutation has no observable effect on the stress response in the brains of mutant mice

In the absence of clear effects of the mutations on global protein synthesis, we hypothesised that the mutations might be affecting neuronal translation in more subtle ways. If E122K impairs translation, it may disrupt ribosome translocation, which could activate cellular stress pathways. We therefore investigated markers of the ribotoxic stress response as an indirect measure in the brain of E122K mice.

Initially, we looked for evidence of XBP1 splicing, as this is the first stage of the unfolded protein response in the ER [33], but RT-PCR analysis revealed no evidence of Xbp1 splicing in the brains of E122K/+ and E122K/E122K mice of either sex at P24 (Figure 2A). We then went on to look at ZAKα [34], a MAPKKK protein associated with the ribosome and which is activated by ribosome stalling, bringing about a ribotoxic response through the activation of the p38 and JNK [35]. However, no change in relative ZAKα phosphorylation (phospho-ZAKα/ ZAKα) was detected in P30 mice homozygous for the E122K mutation, or in heterozygous mice at either 2 or 18 months of age (Figure 2B).

**Figure 2.**
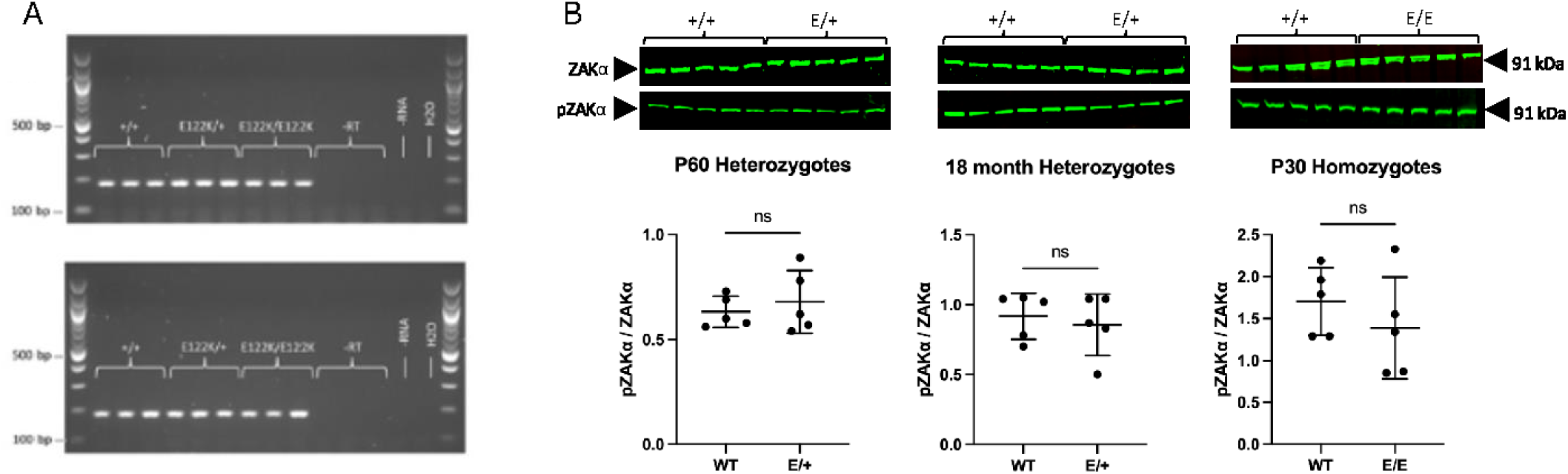
Stress response markers in E122K mouse brains. ***A)*** *Xbp1 RT-PCR products in male (top) and female (bottom) samples. The 500 bp and 100 bp bands of the DNA ladder are labelled. Amplification primers were mXBP1RTspliceF and mXBP1RTspliceR, with an expected amplicon size of 183 bp (unspliced) and 157 bp (spliced). All 18 independent cDNA samples had a single band of 183 bp with no evidence of bands at 157 bp. There was no amplification in -RT, -RNA or water controls*. ***B)*** *Representative immunoblots and quantification of the pZAKá/ZAKá ratio in P60 heterozygous E122K mice, 18-month-old heterozygous E122K mice, and P30 homozygous E122K mice, each compared with age-matched wild-type controls. Brain lysates probed for total ZAKα and phosphorylated ZAKα (Ser165), then quantified in Image Studio Lite and normalised to total protein stain. Five biological replicates per genotype were analysed across two blots [p = 0.685], or P30 homozygous mice [p = 0.365] (two-sample t-test, WT versus other genotypes)*.

### Translatome profiling of hippocampus in mice with the E122K mutation

In the absence of evidence for the E122K mutation resulting in gross changes to protein synthesis, we used a combination of translating ribosome affinity purification (TRAP) and RNA sequencing (RNA-seq) to probe the effects of the mutation on eEF1A2-mediated translation, and to identify mistranslating mRNAs in the mutant mouse brain (Figure 3A). The TRAP methodology uses a bacterial artificial chromosome (BAC) transgenic mouse line, engineered to express a GFP-tagged component of the 60S ribosome, L10a, within specific cell-types [36]. Translating ribosomes can be affinity purified using this tagged ribosomal subunit, enabling the isolation of ribosome-bound mRNA. We used a Snap25-EGFP/Rpl10a line that expresses the GFP-fusion protein only in neurons, allowing us to exclude eEF1A1-expressing glia. We focused on the neuronal translatome in E122K/+ hippocampus, the brain area involved in learning and memory and known to be dependent on protein synthesis to mediate plasticity and memory formation.

**Figure 3.**
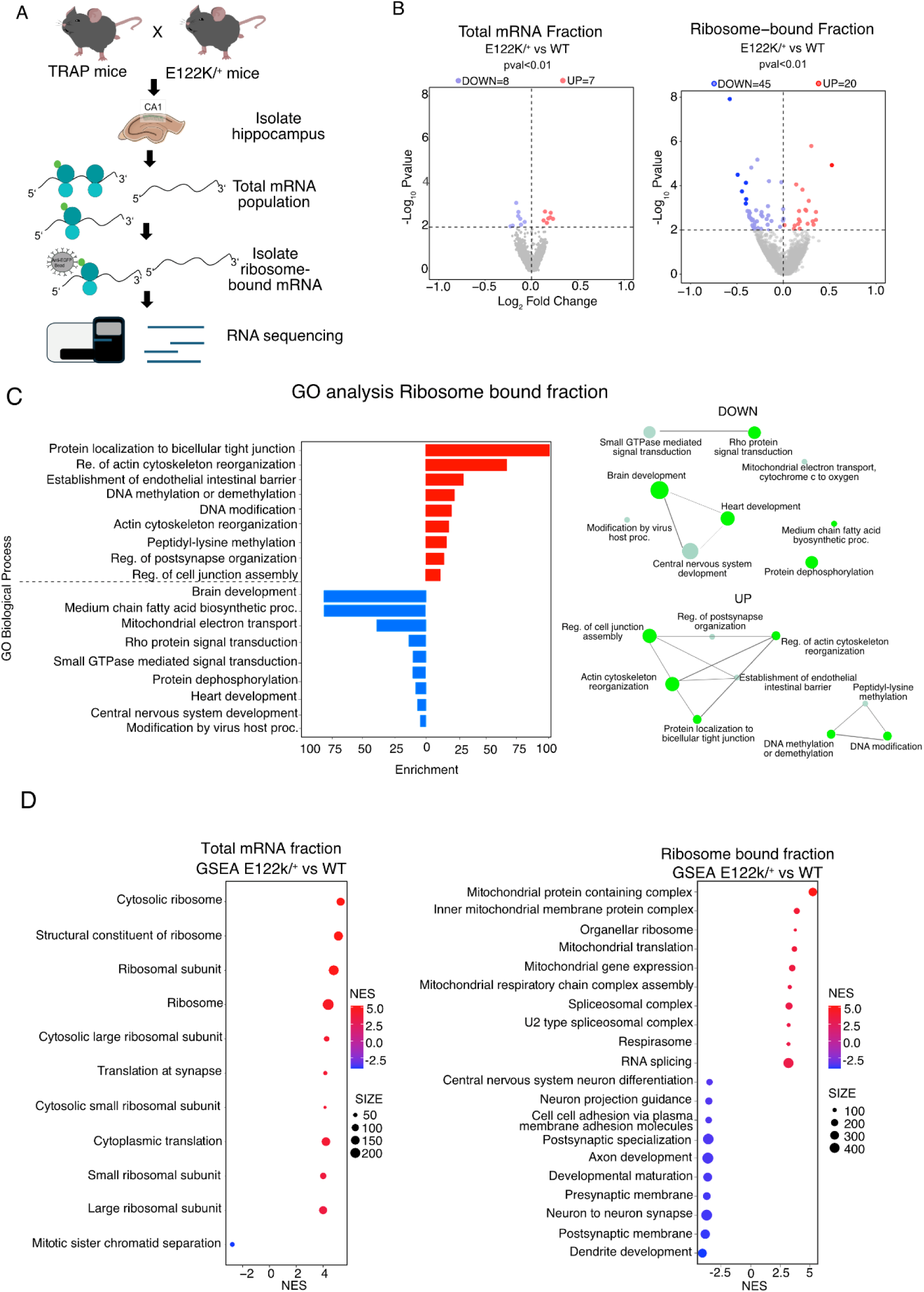
Translatome profiling of hippocampal neurons from E122K/+ mice reveals modest changes of actively translating mRNAs skewed towards downregulation but almost no difference in total mRNA abundance. ***A)*** *Schematic showing the strategy for TRAP-seq*. ***B)*** *Volcano plots for differential expression analysis of E122K/+ vs +/+ hippocampal neurons show changes at threshold p-value <0.01 for the total RNA input (left) and the ribosome-bound mRNA TRAP fraction (right)*. ***C)*** *Left: GO analysis of mistranslated mRNAs for ribosome-bound fraction shows enriched biological processes (BP) amongst the significantly altered genes (p < 0.01) related to actin cytoskeleton reorganization and DNA modification for the upregulated population and Brain development, GTPase signalling, mitochondrial transport for the downregulated set. Most significant enriched terms are shown, selected by an FDR cutoff 0.1 and ranked by Fold Enrichment. Right: Network analysis of the most enriched terms, with an Edge cutoff of 0.3 was performed with Shiny GO*. ***D)*** *Gene set enrichment analysis (GSEA) for gene ontology (GO) terms (biological process, molecular function, cellular component) performed using threshold FDR < 0.05. Depicted are the top 10 upregulated and downregulated categories for the total mRNA fraction (left) and the ribosome-bound fraction (right). Top upregulated pathways for total mRNA fraction include mostly translation machinery related processes while only one downregulated pathway is observed: Mitotic sister chromatid separation pathway. Top upregulated pathways for TRAP dataset include mitochondrial as well as splicing processes while downregulated ones include axon and dendrite development. Each term is represented by dots which are coloured by directionality as expressed by Normalized Enriched Score (NES), and dot size reflects the number of genes in respective terms*.

Hippocampi were extracted from four sets of *Eef1a2*^E122K/+^ and *Eef1a2*^+/+^ littermates at ∼P30, then ribosome-bound RNA was isolated using affinity purification and processed for RNA-sequencing alongside total RNA input fractions. Differential expression analysis between wild-type and mutant samples was performed with DESeq2, identifying 65 differentially expressed genes in the E122K/+ TRAP fraction (Figure 3B) at a significance threshold of p < 0.01. Most of these exhibited a relatively small effect size, with seven genes showing an average log_2_ fold-change greater than 0.4. In comparison, analysis of the total RNA input fraction shows a smaller shift in gene expression. Using the same threshold, 15 genes were identified as significantly different between genotypes, none of which exhibited a log_2_ fold-change greater than 0.4. This suggests that the E122K/+ variant primarily affects translation rather than transcription in the hippocampus. The changes in ribosome-associated transcripts are modest, but are skewed towards down-regulation, with nearly two-thirds of the significantly altered transcripts expressed at lower levels in the mutant samples. However, a similar effect is seen in the input fraction, with 61% of the differential genes downregulated.

Gene ontology (GO) analysis was carried out on all genes with nominal significance (p-value < 0.01). Separate analyses were carried out for upregulated and downregulated transcripts between E122K/+ and +/+ samples in the ribosome-bound fraction only, as the total mRNA fraction was only characterized by small changes. GO analysis in the upregulated population revealed enrichment for genes related to neuronal modelling and cytoskeletal dynamics (Figure 3C), whilst the downregulated population was strongly associated with neuronal development and signalling, as well as metabolic processes. Given the subtle changes observed in both fractions, we also made use of gene set enrichment analysis (GSEA) to identify over-represented classes of genes across the whole dataset. In the TRAP fraction, GSEA revealed the enrichment of GO terms associated with axon and dendrite development in the downregulated population (Figure 3D). The enrichment of these terms suggests that the under-translated mRNAs may most strongly impact neurites rather than the cell body. GO terms relating to mRNA processing, mitochondria, and protein translation were identified as upregulated in the mutant IP fraction indicating an increase in the expression of ribosomal translation machinery, possibly as part of a compensatory response (Figure 3D).

### Long transcripts and SFARI genes are underrepresented in the E122K/+ hippocampal translatome

The translation rate of an mRNA is influenced by transcript length [37], and mutations in ribosomal proteins or in other components of the translational machinery have been found to impact mRNA translation in a length-dependent manner [38, 39]. Additionally, neuronal transcripts are longer on average than non-neuronal transcripts [40, 41], and reduced expression of long genes in the brain has been reported in neurodevelopmental disorders such as Fragile X syndrome [42]. We therefore tested the hypothesis that the E122K eEF1A2 variant exerts a length-dependent effect on translation. Coding sequence (CDS) lengths were obtained from BioMart and binned into four groups (<1 kb, 1-2 kb, 2-4 kb, >4 kb). Analysis of the ribosome-bound dataset revealed a clear length-dependent change that favours shorter transcripts (Figure 4B). A Kruskal–Wallis test followed by Dunn’s post hoc test identified significant differences between all transcript-length groups (adjusted p < 0.01), with progressively lower log₂ fold changes observed as transcript length increased, and the >4 kb group showing significantly more negative log_2_ fold changes than the other groups. This trend was not observed in the total mRNA population (Figure 4A). Transcripts with CDS lengths of 2-4 kb showed significantly higher log₂ fold changes than all other groups, and no significant differences were observed between the remaining groups (adjusted p-value < 0.05). This suggests that the E122K mutation is associated with a length-dependent shift specifically in ribosome-associated transcripts, with a reduced representation of longer mRNAs.

**Figure 4.**
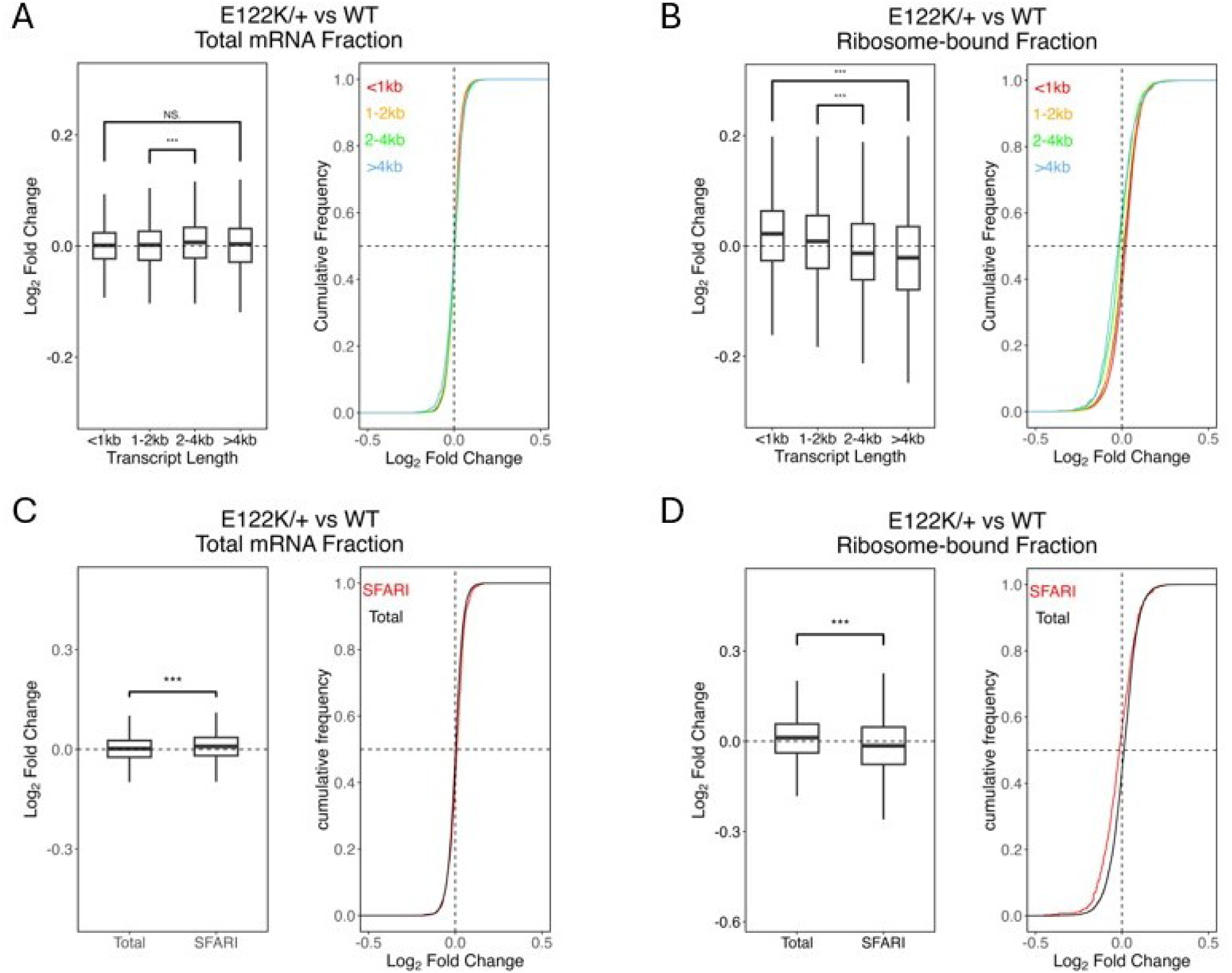
Long transcripts are underrepresented in the E122K/+ hippocampal translatome, as are SFARI genes. ***A-B)*** *Analysis of total mRNA (A) and ribosome-bound mRNA (B) fractions, comparing the log₂ fold change of genes binned by transcript length into four groups: <1 kb, 1–2 kb, 2–4 kb and >4 kb. Transcript lengths obtained from BioMart. Statistical analysis was performed using a Kruskal–Wallis test followed by Dunn’s multiple-comparisons test and Bonferroni correction. **A)** In the total mRNA fraction, transcript length significantly influenced log₂ fold change (Kruskal–Wallis, p < 2.2 × 10⁻¹⁶). Post hoc analysis identified significantly higher log₂ fold changes for transcripts of 2-4 kb compared with transcripts <1 kb, 1–2 kb, and >4 kb, but no significant differences were observed between the remaining groups (adjusted p-value < 0.05). **B)** In the ribosome-bound fraction, longer transcripts showed progressively lower log₂ fold changes than shorter transcripts, with significant differences observed between all transcript-length groups (adjusted p-value < 0.001)*. ***C-D)*** *Subset analysis of SFARI genes (genes in SFARI scoring categories 1–4) compared with the total mRNA and ribosome-bound mRNA populations. Statistical analysis was performed using a Wilcoxon rank-sum test. Whilst significantly higher log₂ fold changes were observed for SFARI genes in the total mRNA fraction (p-value < 0.001), SFARI genes were shifted towards more negative log₂ fold changes in the ribosome-bound fraction (p-value < 0.001), consistent with reduced representation of SFARI transcripts in the ribosome-bound mRNA pool*.

An inherent bias in RNA-seq analyses can lead to the identification of more significant changes in longer transcripts. To determine whether the observed length effect could be driven by this bias, we performed a correlation analysis between CDS length and p-values from the whole dataset, generated from the differential expression analysis. No significant correlation was observed in either the total mRNA fraction (Kendall’s R = -0.0499, p = 0.22) or the ribosome-bound mRNA fraction (Kendall’s R = -0.0314, p = 0.22), indicating that the observed shift is not driven by systemic length bias. To rule out the possibility that the length shift was the result of a different number of transcripts in each CDS bin, we compared CDS lengths between all transcripts and equal-sized subsets (n = 500) of the most significantly up- and down-regulated genes (Supplementary Figure 1B). In the ribosome-bound fraction, the downregulated genes were significantly longer than all transcripts (Wilcoxon, p < 0.0001), whilst the upregulated genes showed no difference (Wilcoxon, p = 0.448). This reflects the CDS length shift observed in the binned analysis in the ribosome-bound population. In the total mRNA dataset, the top 500 most upregulated genes were significantly longer than all transcripts (Wilcoxon, p = 0.00704), whilst the downregulated genes were not significantly different (Wilcoxon, p = 0.0990). This could explain the increased log₂ fold changes observed in the 2-4 kb transcript bin in the total mRNA dataset (Figure 4A).

Longer neuronal transcripts are enriched for genes encoding scaffold proteins, ion channels, and other synaptic components, and are often implicated in neurodevelopmental disorders [39, 40[43]]. We therefore investigated whether genes associated with autism were disproportionately affected in the mutant brain. Analysis of Simons Foundation for Autism Research Initiative (SFARI) genes revealed a significant shift towards downregulation in the E122K ribosome-associated TRAP fraction (Figure 4D). In contrast, a modest but significant shift towards upregulation was observed in the total RNA fraction (Figure 4C).

### Proteomic analysis confirms downregulation of synaptic-specific proteins

Although TRAP-seq enabled us to identify changes in the ribosome-bound population of mRNAs in E122K/+ hippocampal neurons, this analysis alone is unable to provide information about the extent to which these translational changes affect the pool of synthesized proteins, as TRAP-seq cannot inform on ribosome occupancy and thus on translation efficiency. Additionally, diverse studies disentangling the dysregulation in protein synthesis observed in mouse models of Fragile X Syndrome revealed no measurable change in protein abundance in spite of the observed 15-20% increase in protein synthesis [42, 44, 45]. With this in mind, we performed a complementary analysis using label-free liquid chromatography-mass spectrometry (LC-MS) to investigate the proteome of hippocampi from E122K/+ mice of the same age as those used for TRAP-seq. A total of 7,127 proteins were detected, 147 of which exhibited changes with LogFC >| 0.4| and p < 0.01. As with the TRAP analysis, global changes in expression were relatively minor. Unlike the data from the TRAP-seq analysis, there was no detectable skew towards downregulation in the mutant animals, possibly because the LC-MS analysis only detects the most abundant proteins rather than sampling the whole translatome. Of the differentially expressed proteins, 71 were found to be downregulated; among these was eEF1A2, consistent with previous expression analysis [30]. Conversely, 76 proteins were observed to be upregulated (Figure 5A). None of the dysregulated proteins were represented in the TRAP-seq dataset; this divergence between the actively translating mRNAs and the protein abundance could be explained by “buffering” strategies to fine-tune protein synthesis, but it is also important to note that whilst the TRAP-seq specifically represents the translatome of neurons, the proteomics dataset includes contributions from the surrounding neuronal environment, comprising glial/endothelial cells.

**Figure 5.**
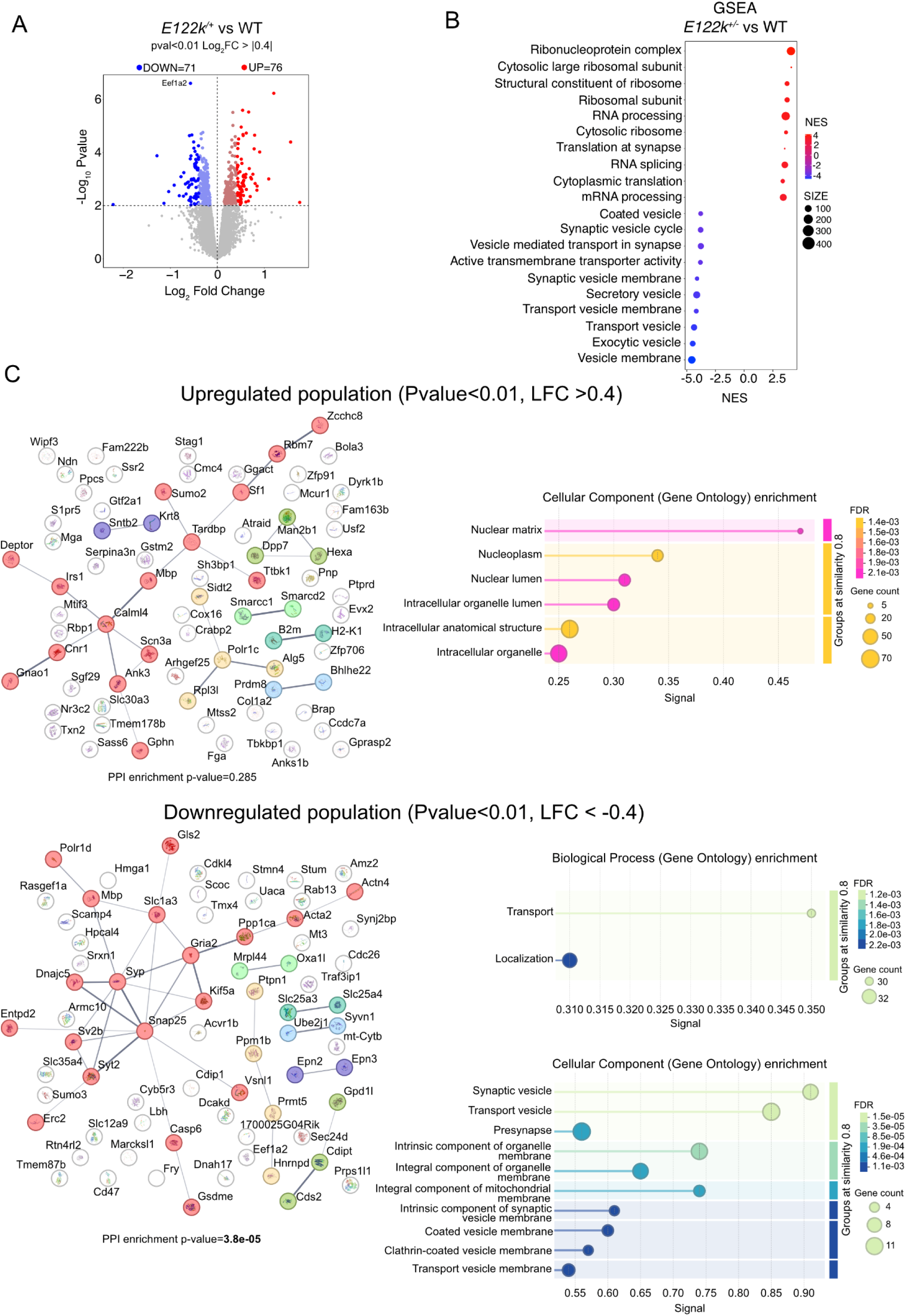
Proteomic analysis of E122K/+ hippocampus reveals downregulation of proteins related to synaptic transmission. ***A)*** *Differential expression analysis of E122K/+ hippocampal proteomics dataset reveals similar protein upregulation (76) and downregulation (71) at the defined thresholds (p-value < 0.01 and log2FoldChange > |0.4|). eEF1A2 protein is confirmed to be significantly downregulated*. ***B)*** *GSEA analysis shows downregulation of synaptic and transport related gene sets in E122K/+ proteomics and upregulation of ribosomal related functions. Plotted are the top 10 significantly upregulated and downregulated categories (FDR<0.05)*. ***C)*** *STRING database was used to generate network plots of protein interactomes respectively for the upregulated and downregulated significantly expressed population. Significant protein interaction (PPI, p-value= 3.8* x 10^-5^*) is observed for the downregulated protein fractions which is enriched for synaptic vesicle and mitochondrial components. The upregulated fraction of proteins enriched for nuclear components does not show significant interaction (PPI, p-value = 0.285)*.

Differentially expressed proteins were interrogated using the STRING database (Figure 5C). A significant protein interaction network was observed for the differentially downregulated proteins (p-value = 3.8 x10^-5^) that was not reproduced in the upregulated population (p-value = 0.285). Moreover, the downregulated fraction displayed a significant enrichment of gene sets associated with synaptic, presynaptic vesicle, and vesicle transport cellular components and biological processes. GSEA revealed upregulation for gene categories including RNA splicing, ribosomal subunit, and translation which were not identified through the GO analysis performed by STRING. The downregulated gene sets enriched by GSEA, on the other hand, confirmed the previously detected downregulation of synaptic- and transport-related gene sets (Figure 5B).

We then combined the results from the GSEA for the proteomics and TRAP-seq datasets. A significant overlap of 75 gene set categories is observed (Supplementary Figure 2). The majority of these were convergently downregulated and included categories relating to synaptic function. Significantly downregulated proteins in the proteomics dataset were better represented in the GSEA downregulated categories in comparison to the downregulated mRNA transcripts in the ribosome-bound dataset. Overall, despite their divergent profiles, both proteomics and TRAP-seq datasets suggest a downregulation of synaptic-related transcripts/proteins.

We next interrogated the proteomics dataset to explore whether a differential representation of SFARI targets was observed, as it was in the TRAP-seq dataset, but no change in protein abundance was detected for SFARI targets (Supplementary Figure 3).

In view of the GSEA analysis of the proteomic dataset suggesting a downregulation of proteins involved in synaptic function, and the fact that this is consistent with the human clinical phenotype, we performed western blot analysis of a selection of key synaptic proteins (SYP, SNAP25 and SYT2; Figure 6A) in tissue extracts from hippocampi of heterozygous mutant mice and wild-type littermates. No significant differences were found between genotypes, possibly because of the relative insensitivity of immunoblotting when detecting relatively subtle changes in expression. However, using synaptosome preparations from the same tissue, we were able to detect a significant downregulation of synaptotagmin 2 (SYT2) in the mutant mice.

**Figure 6:**
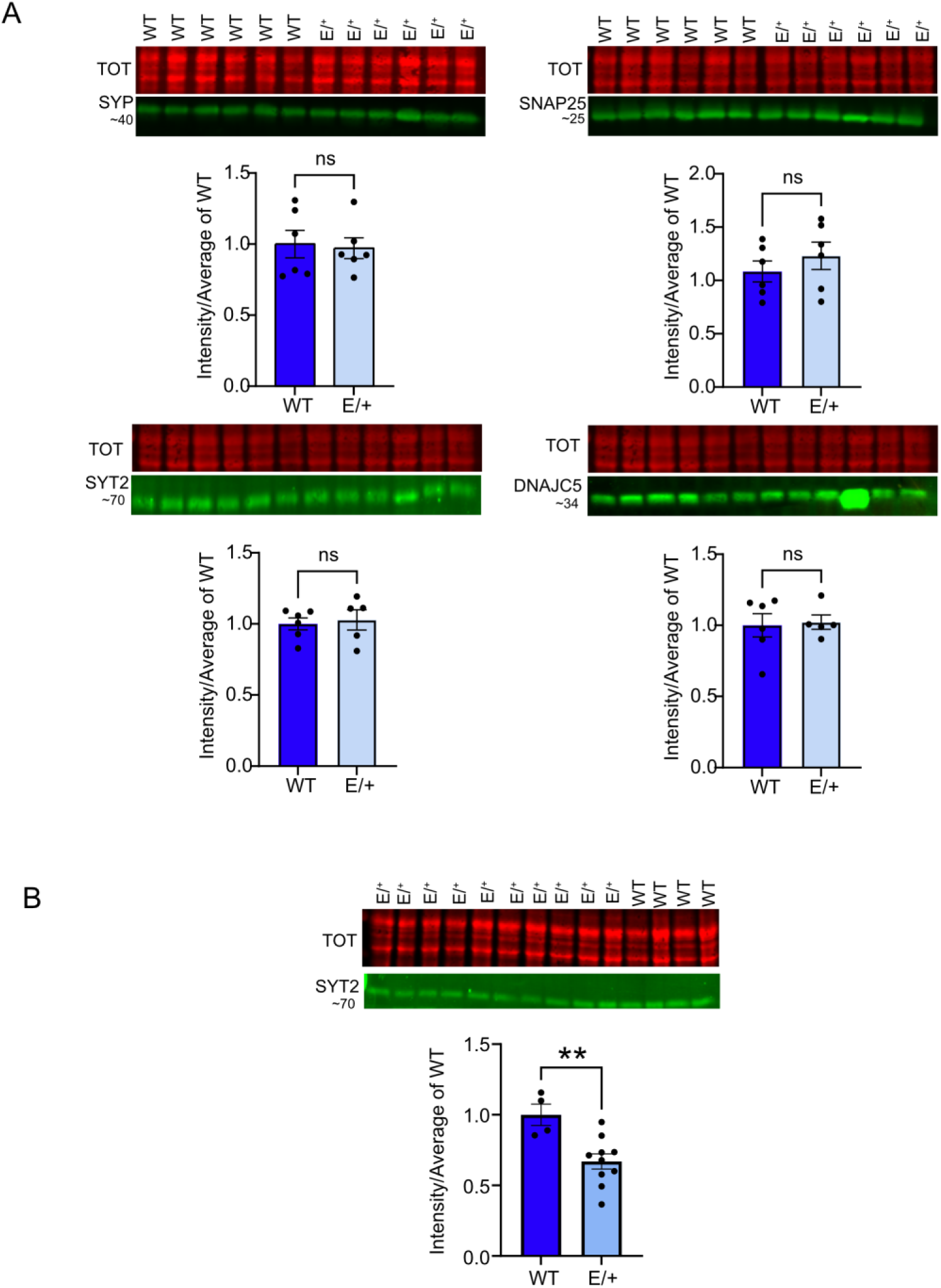
Downregulation of synaptic protein SYT2 is uniquely observed in hippocampal synaptosome enriched fraction but not whole lysate. ***A)*** *Immunoblotting does not confirm downregulation in expression of synaptic proteins in juvenile hippocampus of E122K/+ mice. Four synaptic targets are shown (in order from top left: SYP (5ug-1:50000 dil_Proteintech); top right: SNAP25 (15 ug-1:5000 dil_Merck); below left: SYT2 (15ug-1:1000_SySy); below right: DNAJC5 (15ug-1:5000_Sysy). Replicate E122k/+ 4 for DNAJC5 was an outlier and thus discarded from final analysis. Data are normalized to total protein as well as to average of intensity in WT samples. Unpaired T-test: SYP (p-value = 0.889); SNAP25 (p-value = 0.386); SYT2 (p-value = 0.730); DNAJC5 (p-value = 0.832)*. ***B)*** *Immunoblotting of synaptosome preps from E122K/+ and wild type littermates shows significant downregulation of SYT2 in mutant brain. Synaptosome fractions from E122K/+ and +/+ hippocampi (P28-29) were immunoblotted for SYT2 protein (10ug-1:1000_SySy). Data normalized to total protein as well as to average of intensity in WT samples. Unpaired T-test: SYT2 (p-value = 0.0054)*

## Discussion

Our analysis of gene expression in a mouse model of the most common NDD-causing mutation in *EEF1A2*, E122K, reveals more subtle effects than previously suggested. We set out to replicate the previous study [29] but with additional controls that would allow us to distinguish between two possible mechanisms - either a) a dominant negative effect of mutant eEF1A2 on the function of eEF1A1 (the only form of eEF1A expressed endogenously in HEK cells), or b) enhanced protein synthesis in the cells resulting from transfection with WT eEF1A2, but with reduced enhancement by mutant forms of the protein. Contrary to the previously published results we found no detectable change in translation rates using two different methodologies, not just in primary neurons from the mice with endogenous expression of the E122K mutation, but in transfected HEK cells with any of the three mutations originally reported. The reasons for this discrepancy are unclear, particularly since E122K was reported to reduce protein synthesis to almost half the levels seen in cells with a WT construct.

We went on to examine more subtle effects of the mutation on gene expression in the mutant mice. Further characterization of the hippocampal-neuron specific ribosome-bound fraction by means of TRAP-seq and total transcriptome demonstrated that changes in the translatome are more marked than those in the transcriptome. A modest change in the translation of many transcripts, mostly downregulation, was seen. GO analysis revealed changes in the upregulated translating population related to neuronal modelling and cytoskeletal dynamics, consistent with the reported role for eEF1A2 in the bundling and remodelling of actin within neurons, a process that is key for mediating neurite extension, synaptic formation, and neuronal connectivity [22, 35]. The downregulated translating population suggested that mutant eEF1A2 leads to subtle changes in developmental pathways. Gene set enrichment analysis (GSEA) on the other hand revealed the enrichment of GO terms associated with axon guidance, inhibitory synapses, and the dendritic membrane in the downregulated population, suggesting more of an impact of the mutation in neurites than in the soma.

Analysis of the translatome (but not transcriptome) dataset also revealed a length-dependent change that disfavoured translation of longer transcripts in the mutant hippocampus. Modelling of translation suggests that longer genes, or those with complex UTRs, are translated less efficiently due to the influence of CDS and UTR length on translation initiation, ribosomal recruitment, and ribosome recycling [46, 47]. They therefore require optimal ribosome availability for efficient translation, and some even require tailored *de novo*-produced ribosomes. As neuronal genes tend to be longer and more structurally complex than average [43, 48], changes in ribosome availability may have a more influential regulatory effect on the expression of these proteins. Many of these proteins play a role in synaptic function, dendritic growth, and neuronal maintenance, meaning that altered translation may perturb these pathways, and lead to changes in neural connectivity which are commonly associated with autism and ID [47, 49]. It is well-established that changes in synapse formation and activity can disrupt the balance of excitatory and inhibitory signals in the brain [50, 51]. A decrease in expression of a number of long synaptic genes could therefore impair synapse function and undermine neuronal signaling and communication. The TRAP results thus provide new insights into how the eEF1A2 E122K mutation could translationally impact neuronal function and lead to the associated neurodevelopmental disorders. Interestingly, the reduced expression of long genes in the brain has been reported in other neurodevelopmental disorders, notably Fragile X syndrome and Rett syndrome [42, 52, 53].

The complementary approach of analysing the proteome of hippocampal samples from mutant mice identified changes in expression of different individual genes, but overall pointed to similar perturbations. A significant enrichment was found in the downregulated fraction of proteins for gene sets involved with synaptic function using both STRING database and GSEA analysis.

Follow up analysis of bulk hippocampal samples by western blotting failed to detect the changes seen by mass spectrometry, but when synaptosome preps were made from hippocampus of mutant animals and their WT littermates, significant downregulation of one protein, synaptotagmin 2 (SYT2), was observed. This protein is found in the membrane of synaptic vesicles where it acts as a calcium sensor [54] and has been implicated in motor neuropathy and long-term memory [55, 56]. It is worth noting that although SYT2 shows high levels of co-expression with parvalbumin [57], expression of parvalbumin itself is not significantly altered in the mutant hippocampus. This suggests that the reduction in expression of SYT2 is more specific than a simple reduction in the number of parvalbumin-positive synapses, but this needs to be addressed directly in future experiments.

Many of the changes appear to be focused in axons, dendrites, and synaptic terminals of neuronal cells. This is of interest given that eEF1A1 expression is retained in the axons of mature neurons [16, 58]. In mice, the switch from eEF1A1 to eEF1A2 expression is typically complete by P21, so eEF1A2 would be expected to be exclusively expressed in the cell soma of neurons [12, 13]. It is possible that the changes in the *Eef1a2* E122K hippocampus are primarily a result of a dominant negative effect on eEF1A1, raising a key question concerning the possible treatment and reversibility of the mutant phenotype. Are the observed changes in the TRAP samples a result of mutant eEF1A2-mediated inhibition of eEF1A1 during development, when the two isoforms are co-expressed, or could the mutant eEF1A2 be affecting eEF1A1 activity in axons? Neurons regulate the localization and translation of mRNA in axons locally to control axon guidance, growth, and synaptogenesis. eEF1A is required for protein synthesis, and has also been proposed to mediate mRNA transport, so there is potential for inhibition of eEF1A1 function to compromise synaptic and neuronal connectivity, particularly in light of an increasing awareness of the role of presynaptic and axonal translation [59, 60].

Whilst we detected no evidence for global changes in translation resulting from the presence of mutant eEF1A2, it is clear that there are subtle changes not captured by assays of bulk protein synthesis, and that many of these are likely to impact synaptic function. These changes have the potential to disrupt circuits and result in seizure activity and ultimately in neurodevelopmental disorders. *EEF1A2* is another example of a monogenic cause of neurodevelopmental disorders associated with protein synthesis, and can offer valuable insights into the consequences of even subtle perturbation of protein expression in polarised, long-lived neuronal cells.

## Acknowledgements

We are grateful to the staff of the Bioresearch and Veterinary Facilities at the University of Edinburgh for their expert assistance with the animal work and to the staff of the Proteomics Facility at the IGC for their technical expertise. We are grateful to Alison Meynert and Graeme Grimes of the IGC Bioinformatics core for their support and guidance.

## Materials and Methods

### Mice

For TRAP-seq Snap25-EGFP/Rpl10a from Jackson Labs and *Eef1a2*^E122K/+^ mice were bred on the JAX C57BL/6J background. All experiments were carried out using male littermate mice aged P25-P32 and studied with the experimenter blind to genotype. For Proteomics, *Eef1a2*^E122K/+^ and wild type littermates were used at between P28 and P30, both males and females. Mice were group-housed (5 maximum) in conventional non-environmentally enriched cages with unrestricted food and water access and a 12 h light–dark cycle. Room temperature was maintained in optimal range of 20-24 °C by Air Handling Units (AHU) with humidity at 55%+/- 10%. All procedures were performed in accordance with ARRIVE guidelines and the UK Animal Welfare Act 2006 and were approved by the Animal Welfare and Ethical Review Body at the University of Edinburgh.

### Constructs, cell culture, transfection

Hippocampal and cortical co-cultures were derived from E17.5 mouse embryos. Cells were plated onto glass coverslips coated with poly-D-lysine and 10 μg/ml laminin in an 8-well chamber. Neuronal cultures were incubated in neurobasal media supplemented with 2% B-27 (ThermoFisher Scientific), 0.5 mM L-glutamine (ThermoFisher Scientific), and 1% penicillin/streptomycin antibiotic. Neurons were treated with 1 μM cytosine arabinoside (Ara-C) after 48 hours in order to inhibit non-neuronal cell growth.

HEK-293T cells were cultured in Dulbecco’s Modified Eagles Medium (Gibco) supplemented with 10% (v/v) fetal calf serine (FCS) and 1% penicillin-streptomycin antibiotic. All cell cultures were maintained at 37°C and 5% CO_2_.

### Protein synthesis assays Puromycin assay

HEK293T cells were seeded into 6- or 12-well plates 24 hours prior to transfection with Lipofectamine 3000 (Invitrogen) and either WT or mutant EEF1A2 cDNA within a pcDNA3.1 vector. 48 hours post-transfection cells were treated with 1 µg/ml puromycin for 30 minutes. Cycloheximide was added to control cells for 30 minutes at a working concentration of 50 µg/ml. Cells were washed in in ice-cold phospho-buffered saline (PBS), then harvested with lysis buffer. Cell lysates were incubated on ice for 30 minutes with periodic vortexing, then centrifuged for 10 minutes at 10,000x g and 4°C. Protein concentration was standardised with a bicinchoninic acid (BCA) assay (Pierce), then processed for western blotting with 2x Laemmli buffer, boiling for 5 minutes, then 10% (v/v) Dithiothreitol (DTT). Neurons were treated with 1 µg/ml puromycin for 30 minutes at DIV14, then harvested in the same manner.

### Click-iT AHA assay

AHA assays were carried out with the Click-iT™ AHA Alexa Fluor™ 488 Protein Synthesis HCS Assay Kit (Invitrogen) as per the manufacturer’s instructions. HEK293T cells were seeded in 24-well plates onto sterile coverslips coated with poly-D-lysine. Cells were transfected with Lipofectamine 3000 after 24 hours, then treated with 50 µM Click-iT® L-azidohomoalanine (AHA) reagent in methionine-free RPMI Medium 1640 (Invitrogen) for 30 minutes. Control cells were treated with 50 µg/ml cycloheximide for 30 minutes. Following AHA treatment, cells were incubated in fresh methionine-free media for 10 minutes, washed with ice-cold PBS, then fixed in 3.7% paraformaldehyde (PFA) for 15 minutes. Cells were washed in PBS containing 3% BSA (AHA wash buffer) and permeabilised in 0.5% Triton X-100 for 20 minutes. Cells were washed in AHA wash buffer, then incubated with Click-iT® reaction cocktail for 30 minutes. Following further washes with AHA wash buffer, cells were incubated with Alexa Fluor 597 Phalloidin at 1:500, and Hoechst at 1:1000. Cells were washed twice more with PBS, then mounted in ProLong-GOLD Antifade Mounting Medium (Life Technologies) overnight.

Primary neurons were seeded onto coverslips coated with poly-D-lysine and laminin. At DIV14 cells were incubated with 50 µM AHA reagent for 1 hour. The neurons were then fixed and processed as above. Staining was carried out with rabbit anti Map2 (1:500) at 4°C overnight, and Donkey rabbit Alexa Fluor™ 594 (1:1000) for 1 hour at room temperature. Images were acquired using an EVOS fluorescent microscope at 20x magnification. Cell detection and quantification of Alexa Fluor™ 488 fluorescence intensity in the cell soma were performed in QuPath.

### TRAPseq

Hippocampi were extracted from four sets of *Eef1a2*^E122K/+^ and *Eef1a2*^+/+^ littermates aged ∼30 days and homogenised. Ribosome-bound RNA was isolated using affinity purification using streptavidin/protein L coated Dynabeads bound to anti-GFP antibodies (HtzGFP-19F7 and HtzGFP-19C8) and the Arcturus PicoPure RNA Isolation Kit (Thermo Fisher Scientific). The total RNA input fraction was also collected, and RNA for each sample was quantified and normalised using Quant-it™ RiboGreen RNA Assay Kit (Life Technologies).

### RT-qPCR

Normalised RNA was transcribed into cDNA using Superscript VILO cDNA synthesis kit (Life Technologies), followed by RT-qPCR using Quantitect SYBRgreen qPCR master mix (Qiagen), all according to manufacturer’s instructions. Samples were prepared in triplicate in 96-well reaction plates and run on a StepOne Plus (Life Technologies). For each sample, SNAP25 and GFAP expression levels were normalised to ß-actin and PBIb, and IP samples were normalised to the corresponding inputs to assess TRAP enrichment. The primers used were;

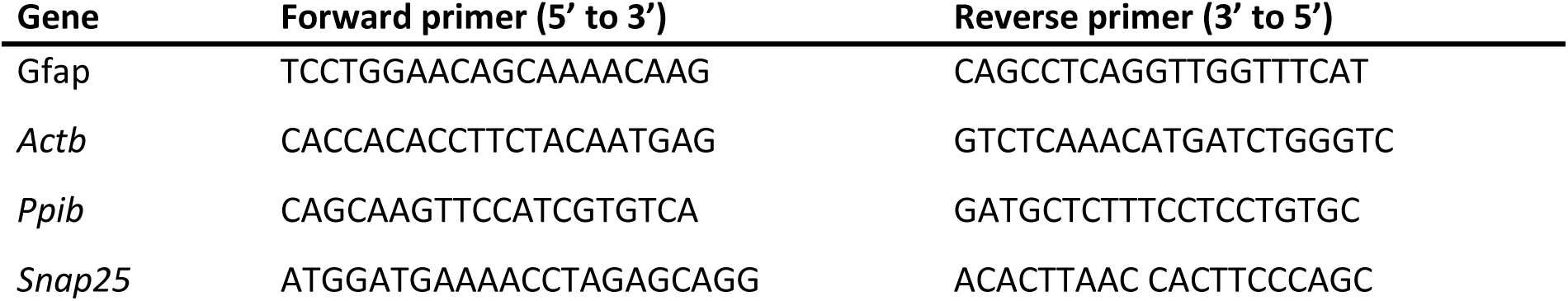

### RNAseq

RNA with an RNA integrity number (RIN) > 7 was then prepared for RNA-seq, using the SMART-Seq® v4 Ultra® Low Input RNA Kit (Takara Bio) and Nextera XT DNA Library Preparation Kit (Illumina) according to manufacturer’s instructions. Paired-end 100bp sequencing was performed on the Illumina NextSeq 2000 platform. All samples generated between 18 and 33 million reads per sample according to SAMtools, and passed quality control checks for per-sequence quality scores, GC content, sequence lengths, duplication, and sequence overrepresentation. Transcript processing, alignment, gene counts, and quality control were performed with nf-core/rnaseq pipeline (1.3.1). Reads were aligned to Ensembl reference genome GRCm39, and gene level counts were quantified across full transcript models. Differential expression analysis was carried out using nf-core/differentialabundance (v1.3.1) and DESeq2 (v1.36.0). Pathway and gene set analysis were performed in R using ShinyGO (v0.741) or the biomaRt and enrichplot R packages.

### Transcript length analysis

Total transcript length, coding sequence (CDS) length, and CDS sequences were obtained from BioMart. The most abundant transcript for each gene was derived from total hippocampal RNA-seq data (TPM values) and used for length analyses. Transcripts were binned into categories by CDS length. Differences in mean CDS length between each category and the overall transcript population were assessed using a Kruskal–Wallis test followed by Dunn’s post hoc test. In addition, a Kolmogorov–Smirnov test was used to compare the distributions of short (<1 kb) and long (>4 kb) transcripts. For analysis, the 500 most upregulated and 500 most downregulated genes by nominal p-value were selected in each dataset.

### Western Blotting

Tissue and cell lysates were prepared in ice-cold RIPA buffer, with protease and in phosphatase inhibitors. Tissue was homogenised using pre-filled DNase/RNase free 1.4 mm ceramic bead mill tubes (Fisher Scientific) using a Precellys-24 sample lyser (Precellys). Lysed samples were centrifuged for 15 minutes at 16,000 x g and 4°C. Protein concentration was determined by BCA assay (Pierce) and lysates were normalised accordingly. Lysates were mixed 1:1 with 2x Laemmli loading buffer, boiled for 5 minutes at 100°C, then 10% (v/v) 1M dithiothreitol was added.

SDS-PAGE was carried out using Mini Gel Tanks (Invitrogen) according to manufacturer’s instructions, using 5 μl Colour Prestained Protein Standard (New England Biolabs) ladder. Proteins were transferred to Immobilon-FL PVDF membranes (Merck) by wet transfer using NuPAGE transfer buffer (Invitrogen). Total protein was stained using REVERT 700 total protein stain (LI-COR), and imaged using a LI-COR Odyssey CLx Imaging System. Blots were blocked using INTERCEPT (TBS) blocking buffer (LI-COR), then incubated with primary antibody solution at 4°C overnight.

Following 4x rinses in TBS-T, secondary antibodies were incubated for 1 hour in the dark then rinsed again. Blots were imaged then analysed using Image Studio Lite, with band intensities normalised either to total protein or a protein loading control.

Mouse brain lysates were prepared as described above from wild-type and mutant mice at P30, P60, and 18 months in order to probe ZAKα levels. ZAKα was detected using a rabbit anti-ZAKα antibody (Proteintech, 28761-1-AP) at 1 in 1000. Phospho-ZAKα (Ser165) was detected using a rabbit anti-phospho-ZAKα (Ser165) antibody (antibodies.com, A51352) at 1 in 1000.

### Xbp1 splicing

RNA extraction: RNA was extracted from mouse brain using the Direct-Zol Miniprep Plus kit (Zymo) according to the manufacturer’s instructions. Off-column DNAse digestion was then performed using the DNA-free™ DNA Removal Kit (Ambion), according to the “routine” version of the manufacturer’s instructions. RNA was then immediately converted to cDNA or stored at -70°C until use. cDNA was synthesised using the High-Capacity cDNA Reverse Transcription kit (Applied Biosystems) according to the manufacturer’s instructions. cDNA synthesis was performed in 96-well plates in 20 μL reactions. 500-2000 μg of template RNA was used per reaction using the programme 25°C for 10 min, 37°C for 2 hours, 85°C for 5 minutes. Control reactions with no RNA (-RNA) or RNA but no reverse transcriptase (-RT) were performed in parallel. Template cDNA was diluted to a concentration of approximately 0.5 ng/ μL for RT-PCR reactions, with a total of ∼0.5 ng of template cDNA per RT-PCR reaction. RT-PCR reactions were performed for each cDNA sample, alongside -RT, -RNA and water controls. Primers were mXBP1RTspliceF GAGAACCAGGAGTTAAGAACACG and mXBP1RTspliceR GAAGATGTTCTGGGGAGGTGAC. The following thermocycler program was used, with a total of 30 cycles: 95°C for 2 min, 30x (95°C for 10s, 60°C for 30s, 72°C for 30s), 72°C for 7 min. To visualise PCR products, 10 μL was mixed with 6x DNA loading buffer and electrophoresed on a 1.5% agarose gel and imaged using a UV transilluminator.

### Synaptosome preparation

Hippocampi from P28-29 mice were dissected in ice and then homogenised in 500 ul of Syn-PER reagent (ThermoFisher Scientific 87793). Homogenate was then centrifuged at 1400x g for 10 minutes at 4°C and 100 ul supernatant afterwards saved as homogenate fraction. The rest was centrifuged at 15000xg for 20 minutes to retrieve the cytosolic fraction as supernatant and synaptosome fraction as pellet. Pellet was resuspended with 200 ul Syn-PER reagent and then BCA assay was performed to quantify all fractions. Samples were then lysed in Laemmli buffer, denatured for 5 minutes at 95 °C and afterwards 0.1M DTT was supplied.

### Mass spectrometry

Hippocampi were dissected from male and female littermate mice and stored at -70°C for future use. Samples were homogenised in RIPA buffer containing protease and phosphatase inhibitors using pre-filled DNase/RNase free 1.4 mm ceramic bead mill tubes (Fisher Scientific) and a Precellys-24 sample lyser (Precellys). Homogenised samples were centrifuged for 15 minutes at 16,000x g and 4°C, then a BCA assay (Pierce) was used to measure protein concentration. 150 μg of each sample was aliquoted and washed in 4x ice-cold acetone, vortexed, and stored at -20°C overnight. Samples were then centrifuged for 10 minutes at 14,000x g, and the supernatant removed. The cell pellet was washed twice more with acetone, each time adding 4x acetone, vortexing briefly to dissolve the pellet, then freezing for 15 minutes at -20°C. The remaining acetone evaporated in a fume hood, then the pellet was suspended in 40 μl of MS Lysis buffer before it became fully dry. Samples were heated to 95°C for 5 minutes, then cooled to room temperature. proteins denatured in urea and alkylated using TCEP and IAA. Peptides were generated by double digestion with Lys-C protease and trypsin, acidified in 100% TFA, and eluted with 50% ACN.

### Proteomics analysis

Peptide abundances from DIA-NN were normalized using variance stabilizing normalization with DEP package 1.26.0 (Bioconductor). Data was first cleaned to remove proteins with missing ID. Filtering was subsequently performed to retain proteins that were expressed in three replicates in at least one condition (low stringency filter). These were then classified as missing not at random (MNAR), while missing values that did not respect these parameters were classified as missing at random (MAR). MAR values were imputed using Bayesian PCA and MNAR values were imputed using quantile regression imputation of left censored data (QRILC). Protein IDs were converted to mouse gene names using biomaRt 2.60.1 (Bioconductor). This yielded 7127 proteins that were considered for downstream analysis.

Differential enrichment analysis was performed using test_diff function from DEP which uses limma and is based on empirical Bayes statistics. Differentially expressed targets were selected based on a significance threshold of p-value <0.01 and Log2FoldChange |0.4|.

### SFARI targets analysis

SFARI transcripts were retrieved from SFARI database (release 07-08-2025). For both datasets, gene ensembl ids were first converted from human to mouse using biomaRt 2.60.1. Expression of both SFARI and epilepsy targets was interrogated for both the significantly expressed population or the total population and compared to total proteomic dataset. Two-side Z test was used to determine differences between dataset distributions.

### STRING analysis

Significantly expressed proteins were submitted to STRING database (version 12.0) to generate independently network graphs for either upregulated or downregulated proteins. Parameters chosen to represent protein network are: network edges displayed by confidence at medium level 0.400 and no additional interactors. Moreover, k-means clustering was applied for each network showing the minimum number of clusters automatically detected. Gene Ontology Enrichment analysis (Cellular Component and Biological Process) was also performed by STRING.

### GSEA and GO analysis

GSEA v4.3.3 was downloaded from (https://www.gsea.msigdb.org/gsea/) and annotated gene sets were retrieved from Molecular Signature Database (v2024.1.Mm) including biological, molecular, and cellular ontology gene sets. GSEA analysis was performed using GSEAPreranked method. After being ranked by fold change, genes from the total proteomic datasets were enriched using “classic” statistics to remove the magnitude bias of ranking metric. Cutoff for identification of genes per gene set was set to minimum 20 and maximum 500 with maximum 1000 permutations. Threshold to retrieve differentially expressed population of gene sets was established at FDR < 0.05. Gene ontology analysis was performed in R using the biomaRt and enrichplot packages. Threshold for retrieval of significantly expressed categories was set at FDR < 0.1 Network analysis of the most enriched terms was performed with Shiny GO (v0.7641) with an Edge cutoff of 0.3.

### Overlap analysis

Overlap analysis was performed with GeneOverlap package 1.40.0.

## Supplementary Material

Sheet 1: Expression data for TRAP-seq total mRNA population E122K/+ vs wild-type.

Sheet 2: Expression data for TRAP-seq ribosome-bound populationE122K/+ vs wild-type.

Sheet 3: Expression data for mass spectrometry E122K/+ vs wild type.

---

**Supplementary Figure 1.**
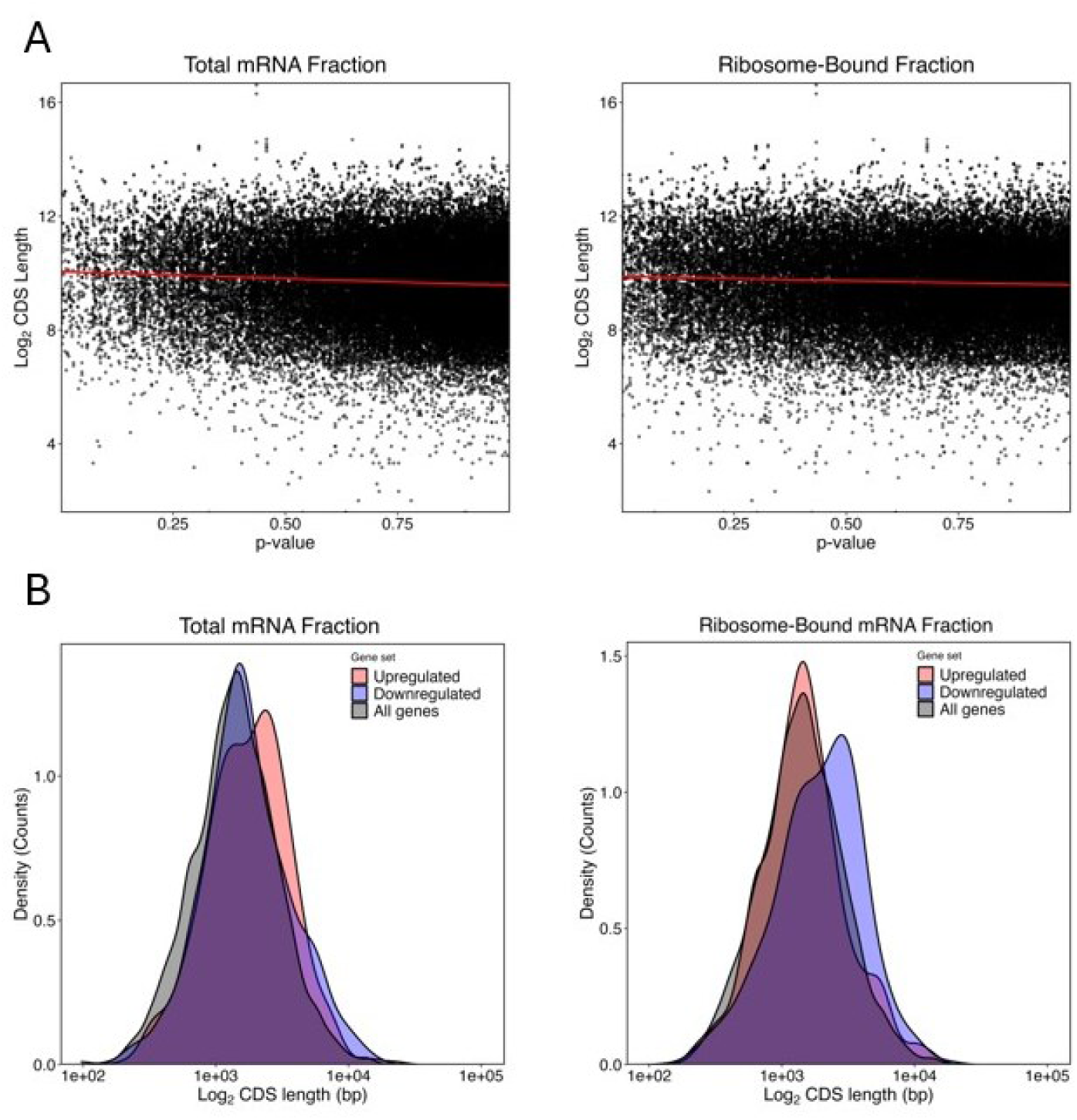
*A. Correlation analysis between CDS length and differential expression p-value, showing that inherent bias in the RNA-seq analysis does not explain the observed difference between long and short transcripts in the Eef1a2*^E122K/+^ *ribosome-bound fraction. Total mRNA fraction (left) Kendall coefficient R = -0.0499, p = 0.22, and ribosome-bound mRNA fraction (right) Kendall coefficient R = -0.0314, p = 0.22*. *B. Density plot comparing the CDS length of the 500 most significantly upregulated (red) and downregulated (blue) genes by p-value, against all genes (black). Transcript data downloaded from BioMart. In the total mRNA dataset, upregulated genes had significantly longer CDS length than all genes (Wilcoxon, p = 0.00704), while downregulated genes showed no significant difference (Wilcoxon, p = 0.0990). In the ribosome-bound fraction, downregulated genes had significantly longer CDS length than all genes (Wilcoxon, p < 0.0001), while upregulated genes showed no significant difference (Wilcoxon, p = 0.448)*.

**Supplementary Figure 2:**
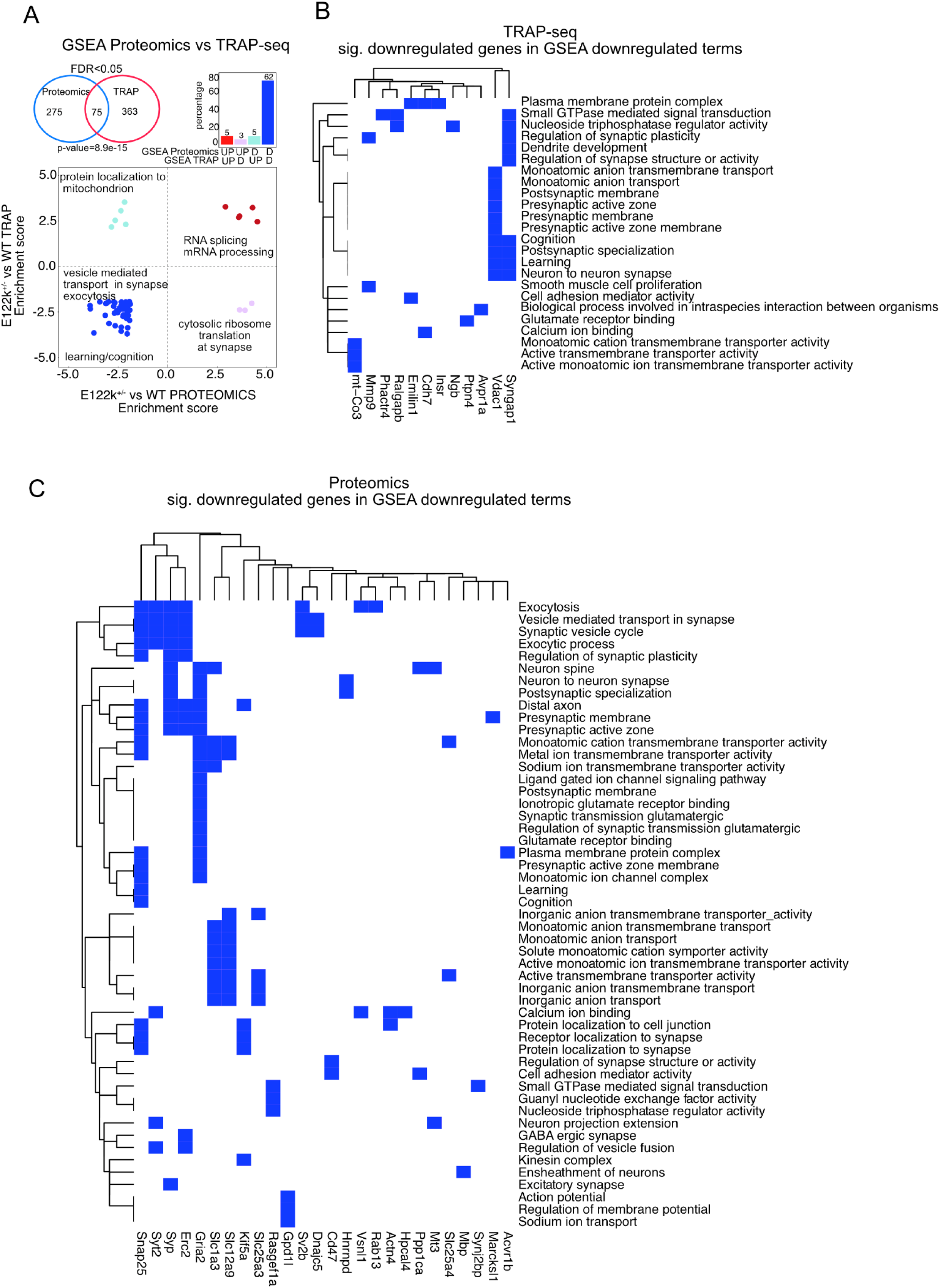
*A. Overlap analysis for GSEA enriched categories in TRAP-seq and Proteomics E122k+/- datasets reveals convergent downregulation of synaptic transmission related pathways. Overlap of GSEA categories (FDR<0.05) from Proteomics and TRAP-seq datasets reveals significant overlap of 75 categories, of which 62 are convergently downregulated and enriched for synaptic related processes (p-value=8.9e-15)*. *B) Significantly expressed genes are shown from TRAP-seq dataset (p-value <0.01, LFC<0) for the converging downregulated GSEA categories as well as from C) Proteomics dataset (p-value <0.01, LFC< -0.4)*.

**Supplementary Figure 3.**
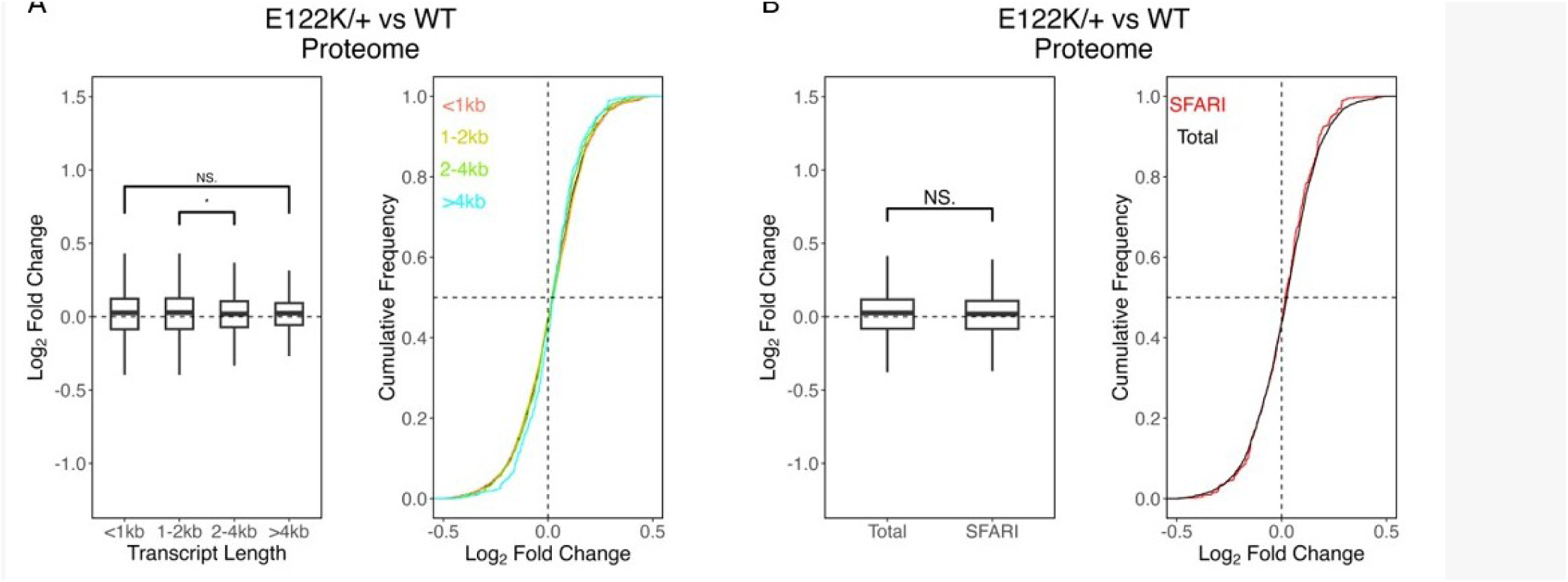
***A***. *Cumulative distribution analysis of E122K/+ and WT proteomic data shows no* significant overall effect of length on log_2_ fold change (Kruskal-Wallis, *p* = 0.181). Post hoc analysis revealed a significantly lower log_2_ fold change in the 2-4 kb group compared with the other groups, but no other significance difference was observed (adjusted p0value < 0.01). **B**) A Wilcoxon rank-sum test found no significant difference comparing the SFARI gene subset with the total population (p = 0.181).

